# Bipartite influence of abscisic acid on xylem differentiation trajectories is dependent on distinct VND transcription factors in Arabidopsis

**DOI:** 10.1101/2020.09.28.313189

**Authors:** Prashanth Ramachandran, Frauke Augstein, Shamik Mazumdar, Thanh Van Nguyen, Elena A. Minina, Charles W. Melnyk, Annelie Carlsbecker

**Affiliations:** Department of Organismal Biology, Physiological Botany, Linnean Centre for Plant Biology, Uppsala University, Ullsv. 24E, SE-756 51, Uppsala, Sweden; Department of Plant Biology, Linnean Center for Plant Biology, Swedish University of Agricultural Sciences, Ullsv. 24E, SE-756 51, Uppsala, Sweden; Department of Molecular Sciences, Linnean Center for Plant Biology, Swedish University of Agricultural Sciences, Ullsv. 24E, SE-756 51, Uppsala, Sweden

**Keywords:** ABA, *Arabidopsis thaliana*, plasticity, VND, water deficiency, xylem differentiation

## Abstract

Plants display a remarkable ability to adjust their growth and development to changes in environmental conditions, such as reduction in water availability. This high degree of plasticity is apparent not only as altered root and shoot growth rates, but also as changes to tissue patterning and cell morphology [1,2]. We have previously shown that *Arabidopsis thaliana* root xylem displays plastic developmental responses to limited water availability, mediated by non-cell autonomous action of abscisic acid, ABA [2]. Here, we show through analyses of ABA response reporters and tissue specific suppression of ABA signalling that xylem cells act as primary signalling centres for mediation of changes to both xylem cell fate and differentiation rate revealing a cell autonomous control of xylem development by ABA. Transcriptomic changes in response to ABA showed that members of the VASCULAR RELATED NAC DOMAIN (VND) transcription factor family are rapidly activated. Molecular and genetic analyses revealed that the two aspects of xylem developmental changes, cell fate and differentiation rate, are dependent on distinct members of this transcription factor family. Thus, this study provides insights into how different aspects of developmental plasticity can be interlinked, yet genetically independent of each other. Moreover, similarities in phenotypic and molecular responses to ABA in diverse species indicate an evolutionary conservation of the ABA-xylem development regulatory network among eudicots. Hence, this study gives molecular insights on how environmental stress promotes anatomical plasticity to key plant traits with potential relevance for water use optimization and adaptation to drought conditions.

## Results and Discussion

### ABA affects both xylem cell fate and differentiation rate in Arabidopsis roots

We and others have previously shown that water limiting condition triggers the initiation of multiple protoxylem-like cells with spiral secondary cell walls (SCW) (Fig. 1A, B, C, E, S1B; [2,3]). This effect is partly dependent on ABA signaling within the endodermal cell layer, surrounding the vascular stele, resulting in enhanced levels of microRNA165. This miRNA acts as a non-cell autonomous signal suppressing target HOMEODOMAIN-LEUCINE ZIPPER class III (HD-ZIP III) transcription factors within the stele, thus promoting protoxylem over metaxylem cell fate [4,5]. Interestingly, a recent study showed that in certain Arabidopsis ecotypes metaxylem form closer to the root tip, and this may lead to enhance hydraulic conductance and better survival under water limiting conditions in certain Arabidopsis ecotypes [6]. We, therefore, hypothesized that other ecotypes may achieve a similar phenotype by enhancing xylem differentiation rates, and that this could be a plastic developmental response to water limiting conditions. To assess this, we first analyzed the distance at which secondary cell wall (SCW) lignification was observed in the Colombia (Col-0) ecotype after treatment with 1µM ABA. We have previously shown that this treatment can be used as a proxy for water limiting conditions without the negative impact on root growth, and that may confound analysis of xylem differentiation rate (Fig.S1A; [2]). A 48h ABA treatment caused protoxylem cells occupying the outer position (*px* position) of the xylem axis traversing the Arabidopsis root (Fig. 1A, B) to differentiate slightly closer to the root tip (*px* mock at 1264±139µm (SD) *vs.* ABA at 1060±240 µm), whereas the metaxylem cells next to these cells (*outer metaxylem, omx*, position), differentiated significantly closer to the root tip (*omx* mock 2950±374µm vs ABA 1912±393µm; Fig.1F, G). However, as previously described [2] the cells in the *omx* position frequently underwent a change in fate upon ABA treatment resulting in xylem vessels with reticulate or even protoxylem-like spiral SCW rather than the pitted morphology usually observed in Arabidopsis primary root metaxylem (Fig. 1C, E, S1B). Because protoxylem cells normally differentiate first and thereby closer to the root tip (hence ‘proto’), the observed earlier differentiation in the *omx* position may be coupled to the change in fate. In contrast to the *omx*-cells, we found that the *inner metaxylem, imx*, cells never formed reticulate or spiral SCW upon 1µM ABA treatment (Fig. 1C), suggesting that fate change did not occur in these cells. Therefore, we monitored the cells occupying the *imx* position that normally differentiate late and at a considerable distance from the tip (generally >7 mm from the tip in 5-day old seedlings). Despite the absence of any change in SCW morphology, a 48h treatment with 1µM ABA resulted in 94% of the roots displaying differentiated *imx* at 7 mm from the root tip (Fig.1H) compared to 0% in mock-treated plants, suggesting that ABA promotes metaxylem differentiation rate independent of its effect on xylem morphology or root growth. Furthermore, transferring back to mock conditions restored both normal xylem morphology and differentiation rate within 48h (Fig.S1C,D,E), corroborating the notion of considerable developmental plasticity in the formation of plant xylem.

**Figure 1:**
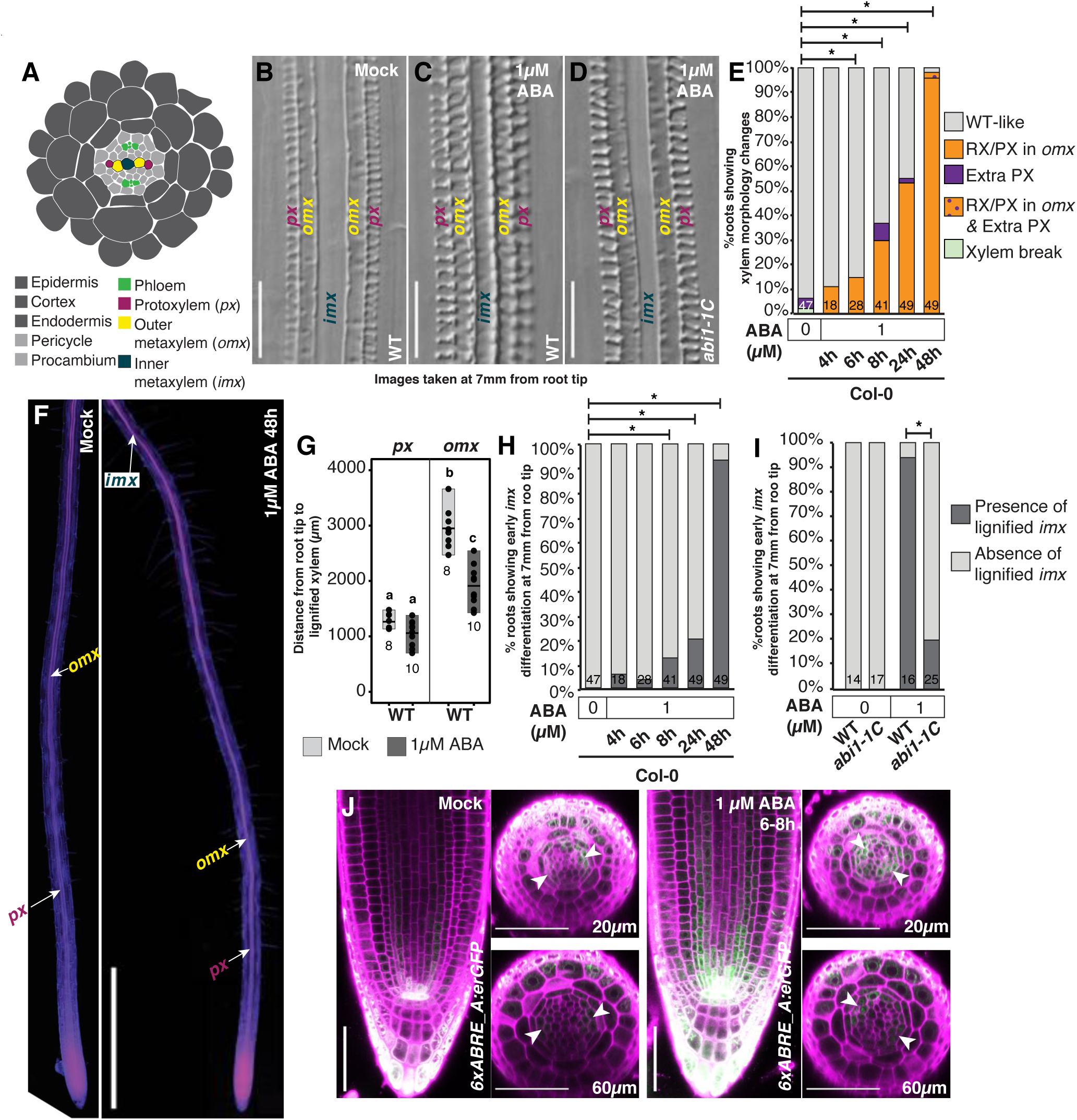
ABA affects both xylem differentiation fate and rate in Arabidopsis roots. **(A)** Cartoon of a cross section of the Arabidopsis root showing different cell types highlighting the different positions in the xylem axis, *px, omx* and *imx*. **(B-D)** Differential Interference Contrast (DIC) images of the xylem pattern at 7mm from the root tip after ABA treatment in WT (B, C) and *abi1-1C* (D). **(E)** Temporal analysis of xylem morphology changes in WT roots after 1µM ABA treatment for 4h, 6h, 8h, 24h and 48h. For the 4h, 6h and 8h ABA treatments, the total treatment time before root xylem analysis was 24h. **(F)** Mock and ABA treated WT roots double stained with basic fuchsin (magenta) and calcofluor white (blue). Pink, yellow and blue-green text with white arrows indicate the first occurrence of a fuchsin stained xylem vessel in the *px, omx* and *imx* positions respectively. Under mock conditions, differentiated xylem in the *imx* position was detected at a distance > 7mm from the root tip, and not included within the imaged region of the root. **(G)** Quantification of distances from the root tip to the presence of first lignified vessel in *px* (left) and *omx* (right) positions. In the *px* position, spiral walled xylem vessels were present under both mock and ABA treatments, in *omx* position mock treated roots had pitted SCW while ABA treated roots had either pitted/reticulate/spiral SCW. The difference in morphology of vessels in *omx* position was not considered for the quantification of distances. Black filled dots represent measurements from individual roots. **(H)** Quantification of frequency of roots showing presence/absence of a lignified xylem vessel at the *imx* position in the lower 7mm of the root from the root tip, after 1µM ABA treatment. The treatment times are as in (E). **(I).** Frequency of early *imx* differentiation measured as in (H) in WT and *abi1-1C* after 48h 1µM ABA treatment. **(J).** Confocal micrographs showing the ABA response domains in the root apical meristem visualized using *6xABRE_A:erGFP* after mock or 1µM ABA treatments. Radial optical sections were taken at 20µm and 60µm shootward of the quiescent center (QC). Magenta: Propidium iodide, Green: GFP and white arrow heads: xylem axis. RX, reticulate xylem; PX, protoxylem; *px*, protoxylem position; *omx*, outer metaxylem position; *imx*, inner metaxylem position. Scale bars: 50µm in B and D, 1mm in F. Statistics: * in E, H and I represent P<0.05, Fisher’s Exact Test. In G, *a,b,c* represent groups with significant differences, one way ANOVA with Tukey’s post-hoc testing (P<0.05). Numbers at the bottom of the bars in E, G, H and I represent the number of roots analyzed.

The importance for canonical PYL/PYR receptor-PPC2 mediated ABA signaling in affecting xylem differentiation was evident by the ability for the dominant *ABA INSENSITIVE 1* mutant (*abi1-1*), in which ABA signaling is suppressed even in the presence of ABA [7,8], to strongly reduce the effects of ABA treatment on early *imx* differentiation (20% in *abi1-1* vs 94% in wildtype; Fig. 1D,I), and on xylem fate change in *omx* (Fig.S1F; [2]). However, while repression of ABA signaling specifically in the endodermis (using *pSCR*::abi1-1) could significantly suppress the *omx* fate change (Fig. S1G; [2]), it had less effect on the enhanced differentiation (65% vs 100%; Fig. S1H), suggesting that ABA signaling in other cell types contribute to these dynamic xylem responses. To determine the cell-types in which ABA response occurs we used available synthetic ABA responsive reporters with tandem ABRE element repeats of two ABA responsive genes, *ABI1* and *RAB18* (*p6XABRE_A:GFPer* and *p6XABRE_R:GFPer*) [9]. Under mock conditions, both reporters revealed weak expression in the epidermal cells and the *p6XABRE_A:GFPer* line also displayed weak expression in the endodermal, xylem pole pericycle and protoxylem precursor cells suggesting ABA signaling occurs in these tissues under non-stressed conditions (Fig. 1J, S1I). After 6-8h treatment with 1µM ABA, the signal intensity of both reporters increased in these tissues, and, within the stele, an ABA response maximum was observed in the xylem precursor cells, especially in the *px* position (Fig.1J,S1I). Next, we simulated water stress by growing plants on media overlaid with 550g/l polyethylene glycol (PEG; Fig. S1J) which generated a negative water potential of −1.2 MPa and thus limited water availability. Here, we observed a similar but stronger ABA response suggesting that exogenous ABA treatment could recapitulate the cell-specific ABA response observed during water deprivation.

### ABA signalling within the xylem activates VND transcription factors

The ABA response profile prompted us to investigate the importance of ABA signaling in different tissue domains for xylem differentiation. We analyzed F1 progeny of the *UAS:abi1-1* [10] line crossed with transactivation lines expressing in the xylem axis, *J1721*, procambium, *Q0990*, or ground tissue, *J0571*, upon ABA treatment (Fig.2A, Fig. S2A). We found that the *abi1-1* xylem driver line could efficiently suppress ABA’s effects on both xylem differentiation rate and fate (Fig. 2B,C, Fig. S2B). Neither the procambium nor the ground tissue *abi1-1* driver lines could suppress xylem differentiation rate, but consistent with our previous observations the ground tissue line could partially suppress the xylem fate changes (Fig. 2B,C; S2B;[2]). Furthermore, while the mock treated ground tissue line occasionally displayed discontinuous metaxylem [2], neither of the stele driver lines displayed altered xylem differentiation under mock conditions (Fig. S2B,C). These results suggest that basal ABA signaling in the stele is not critical for xylem formation *per se*, but that signaling within the xylem cells themselves is essential for plastic changes in both xylem differentiation rate and fate upon conditions causing elevated ABA levels.

**Figure 2:**
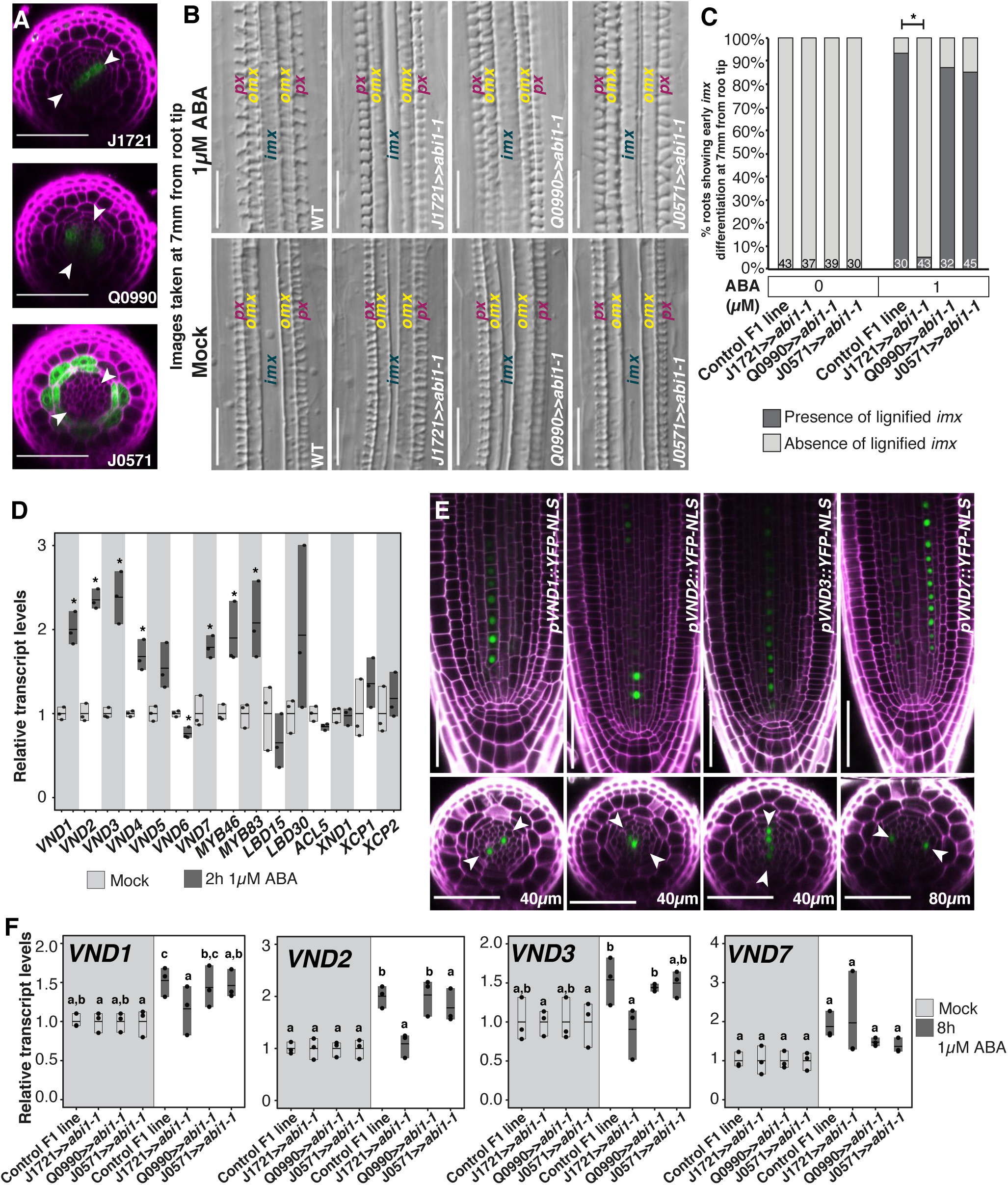
ABA signalling within the xylem activates VND transcription factors. **(A)** Radial optical sections representing the activity domains of the J0571, Q0990 and J1721 GAL4-enhancer trap lines imaged in F1 individuals from crosses with *UAS:abi1-1*. Arrowheads mark the xylem axis. **(B)** DIC images of the xylem pattern in 48h 1µM ABA and mock treated *abi1-1* transactivation lines. **(C)** Quantification of early xylem differentiation in *imx* after mock and 1µM ABA treatment in different *abi1-1* transactivation lines. **(D)** Relative transcript levels of xylem development genes in 1mm WT root tips after 2h of 1µM ABA treatment using qRT-PCR. **(E)** Confocal images showing the transcription domains of *VND1, VND2, VND3* and *VND7* in the longitudinal and radial planes. **(F)**. Relative transcript levels *of VND1, VND2, VND3* and *VND7* after 8h ABA treatment in whole roots of F1 seedlings from cross between *UAS:abi1-1* and indicated GAL4 enhancer trap line quantified using qRT-PCR. *omx*, outer metaxylem position; *imx*, inner metaxylem position. Scale bars: 50µm in A, B, and E. Statistics: * in C, represent P<0.05, Fisher’s Exact test; * in D represent P<0.05, two tailed Student’s t-test. In F, *a,b,c* represent groups with significant differences, one way ANOVA with Tukey’s post-hoc testing (P<0.05). Numbers at the bottom of the bars in C represent the number of individuals analyzed. In D and F, all values are normalized to the average of respective mock treated samples.

Consistent with the observation that ABA signaling in xylem cells is important for their developmental trajectories, transcriptome analysis of 8h ABA-treated roots revealed differential expression of more than 200 genes that were previously assigned to a xylem expressed cluster in a single cell RNA sequencing study on roots [11] (Fig. S3ED, Table S1). Among genes responding to ABA we found down regulation of HD-ZIP III genes (*ATHB8* and *REV*), as expected from previous studies [2], but also upregulation of multiple xylem differentiation regulators along with cellulose and lignin-biosynthesis genes (Table S1). We identified the closely related *VND2, VND3* and *VND7* transcription factors [12–15] among upregulated xylem developmental regulators, suggesting that several VND-gene family members are ABA regulated. To test this observation further, we performed qRT-PCR after 2, 4 and 8h 1µM ABA treatment in root tips for all VND factors along with a number of other xylem differentiation regulators (Fig.2D, S2D,E). We found that 2h of 1µM ABA treatment was sufficient to significantly upregulate *VND1, VND2, VND3, VND4*, and *VND7*, while longer treatment times induced also *VND5* whereas *VND6*, previously found to promote metaxylem differentiation [12], was not upregulated, even after treatment with higher concentrations of ABA (Fig.2D,S2F).

Analysis of transcriptional reporter lines revealed that *VND1, VND2* and *VND3* express in immature xylem cells within the meristem, with *VND1* expression restricted to *omx* cells, *VND2* to all metaxylem precursor cells (*omx* and *imx*), while *VND3* expression was observed in *px, omx* and *imx* cells (Fig.2E). *VND3* expression extended into the differentiation zone, while *VND1* and *VND2* were restricted to the meristem (Fig.2E,S2G). *VND7* expressed specifically in the protoxylem precursors within the meristem and continued beyond the meristem primarily within these cell lineages [5] (Fig.2E,S2G). Significantly enhanced levels but no change in expression pattern of *VND3::NLS-YFP* was detected after 1µM ABA treatment for 6-8h (Fig.S2H,I).Consistent with the notion that ABA signaling within the xylem axis is required for their elevated levels, the xylem J1721>>abi1-1 driver could suppress the activation of *VND1, VND2* and *VND3* by 1µM ABA (Fig. 2F). Taken together, these data show that VND gene expression rapidly and specifically increases within the xylem precursor cells upon increase in ABA levels.

### VNDs regulate plasticity in xylem fate and differentiation rate

To test the requirement for VND transcription factors for the ABA-induced xylem developmental changes we analyzed *vnd* mutant xylem phenotypes after ABA treatment and after growth under water limiting conditions. Under mock conditions, *vnd1, vnd2, vnd3* and *vnd7* single and most double mutant combinations displayed a wildtype-like xylem pattern (Fig.S3A). However, the *vnd2vnd3 (vnd2,3)* and *vnd1vnd2vnd3 (vnd1,2,3)* mutants displayed discontinuous metaxylem strands (Fig. 3A,B,D) and in *vnd1,2,3* the metaxylem strand in one of the *omx* and in the *imx* position frequently failed to differentiate entirely (Fig.3D). Upon ABA treatment *vnd2,3* and *vnd1,2,3*, and to some extent also *vnd2* and *vnd3* did not display early xylem differentiation in the *imx* position seen in ABA-treated wildtype plants (Fig.3A,C; Fig.S3B). Exposure of wildtype plants to water limiting conditions, achieved by growth on PEG-overlaid media, resulted in early xylem differentiation in the *imx* position, but also reduced root growth, rendering it difficult to measure the extent of the early xylem formation. However, importantly, in *vnd1vnd3, vnd2,3* and *vnd1,2,3* mutants the early xylem phenotype was suppressed, although root growth was affected similarly as in wildtype (Fig. 3D,E, S3C,D). Hence, these VND factors are required to promote early xylem differentiation in the *imx* position, upon ABA signaling and under water limiting conditions.

**Figure 3:**
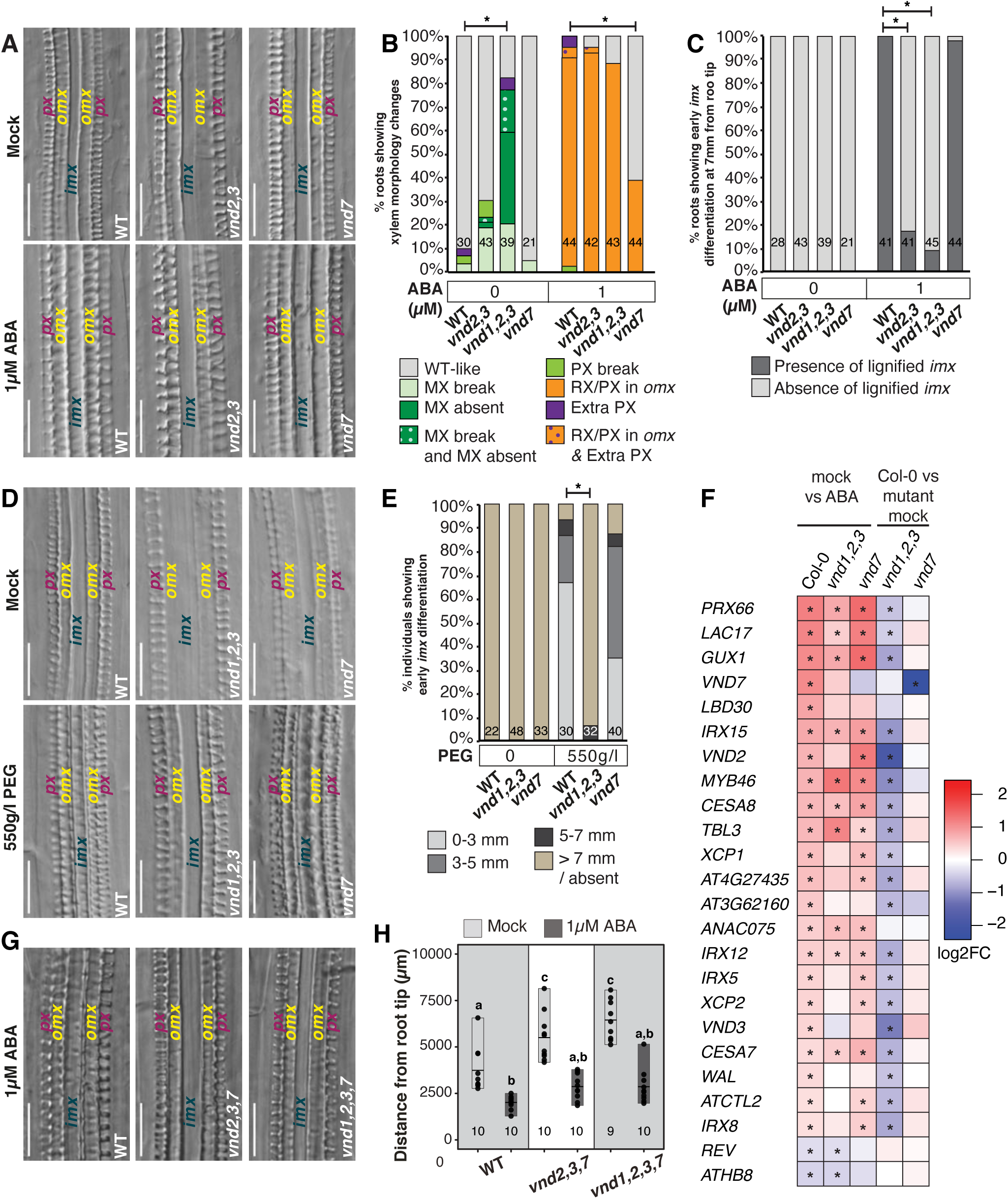
VNDs regulate plasticity of xylem fate and differentiation rate. **(A)**. Representative DIC images of mock and ABA treated wildtype (WT), *vnd2 vnd3* (*vnd2,3*) and *vnd7* root xylem taken at 7mm from the root tip. **(B-C)** Quantification of xylem morphology (B) and differentiation at 7 mm in *inner metaxylem position* (*imx*) (C) changes in *vnd2,3, vnd1 vnd2 vnd3* (*vnd1,2,3*) and *vnd7* mutants. **(D)** Representative DIC images of mock and polyethylene glycol (PEG) treated WT, *vnd1,2,3* and *vnd7* root xylem. **(E)** Quantification of distances at which the first signs of differentiated *imx* was detected in WT, *vnd1,2,3* and *vnd7* roots subjected to mock or PEG treatments. **(F)** Heatmap showing RNA sequencing results displaying the response of a subset of xylem related genes upon ABA and in WT, *vnd1,2,3* and *vnd7* mutants. **(G)** DIC images showing xylem pattern in WT, *vnd2,3,7* and *vnd1 vnd2 vnd3 vnd7* (*vnd1,2,3,7*) mutants after 1µM ABA treatment. **(H)** Quantification of distances from the root tip to the first sign of lignified xylem in *omx* position for WT, *vnd2 vnd3 vnd7* (*vnd2,3,7*) and *vnd1,2,3,7*. Scale bars are 50µm in A, D and G. Statistics: * in B, C, E represent P<0.05, Fisher’s Exact test; * in F represents Padj<0.05; in H, *a,b,c* represent groups with significant differences, one way ANOVA with Tukey’s post-hoc testing (P<0.05). Numbers at the bottom of the bars in B, C, E and H represent the number of roots analyzed.

Despite having a role in *imx* differentiation rate, *vnd2,3* and *vnd1,2,3* displayed protoxylem-like or reticulate xylem morphology in the *omx* position as well as early xylem differentiation in this position upon ABA treatment (Fig.3A,B S3F,G). Instead, we found a requirement for *VND7* in controlling the *omx* fate change because the shift to reticulate xylem and protoxylem in the *omx* position upon ABA treatment was partially suppressed in the *vnd7* mutant (Fig.3A, B). This is in line with the expansion in *VND7* expression into metaxylem cells upon high concentration of ABA, previously noted by Bloch et al [16]. The *vnd7* mutant, however, responded with early *omx* and *imx* differentiation similar to wildtype (Fig.3C, S3F,G). Hence, treating the *vnd7* mutant with ABA revealed a previously uncharacterized necessity for VND7 in the change in xylem cell fate from metaxylem towards protoxylem-like cells. Furthermore, our results indicate that ABA’s effect on xylem differentiation rate can be genetically separated from its effect on xylem cell fate change via the activation of distinct VND genes.

To further dissect the impact of these VNDs in xylem developmental plasticity upon ABA treatment we analyzed the transcriptomic effects in the *vnd1,2,3* and *vnd7* mutants (Table S1.). Confirming the importance of these factors for xylem differentiation, ABA’s induction of xylem cell death regulators, *XCP1* and *XCP2*, secondary cell wall related genes such as *WAL, IRX5, IRX8, AtCTL2* were suppressed in *vnd1,2,3*. In total, 95 of the 223 ABA-induced xylem enriched genes, were dependent on the VNDs (Fig. S3E). We also found SCW related genes, such as *IRX12, IRX15, TBL3, CESA7* and *CESA8*, as well as *MYB46* and *ANAC075* [17– 19], that were upregulated upon ABA treatment in wildtype and *vnd1,2,3*, suggesting the presence of other upstream activators of these factors in response to ABA. Since *MYB46* is down stream of VND7 [20] and VND1, VND2, VND3 and ANAC075 are upstream regulators of *VND7* [13], we reasoned that all four VNDs might act redundantly with each other, in particular in *omx* differentiation, and we therefore generated a *vnd1vnd2vnd3vnd7* mutant. In this mutant, the fate change upon ABA treatment in the *omx* positions was prevented, as well as the premature differentiation in the *imx* position (Fig. 3G,S3H), showing the additivity of the two phenotypes. However, although the cells in the *omx* position maintained a metaxylem morphology with pitted cell walls, they could still respond to ABA with faster differentiation similar to wildtype (Fig, 3G, H). Hence, factors other than VND1, VND2, VND3 and VND7 contribute to the early differentiation in this position or they act redundantly with the VNDs in controlling xylem differentiation rate.

### ABA induces ectopic lignification in cotyledons

Analysis of the genes responding to ABA identified a considerable overlap with genes induced upon tracheary element trans-differentiation in Arabidopsis cell suspension cultures [12]. We therefore asked if ABA could induce trans-differentiation of mesophyll to xylem cells in cotyledons, as previously seen to occur upon treatment with a cocktail consisting of bikinin, an inhibitor of GSK3 kinases involved in brassinosteriod signaling, along with auxin and cytokinin [21]. We substituted bikinin for ABA and, strikingly, we found ectopic lignification occurring in cells encompassing 50-70% of the cotyledon area (excluding the normal venation) (Fig.4A, B, S4B,C), an effect suppressed in the *abi1-1* mutant to 30% (Fig. 4B). However, these ectopically lignified cells did not form properly patterned SCW, as observed upon bikinin treatment [21], suggesting that ABA treatment could not induce the full differentiation program of tracheary element cells during the trans-differentiation process. Nonetheless, the formation of ectopic lignification was suppressed in the *vnd1,2,3* and *vnd7* mutants (Fig. 4A,B), suggesting that the ectopic formation of ABA induced lignified cell walls is at least partly mediated by the VND1, 2, 3, and 7 transcription factors.

**Figure 4:**
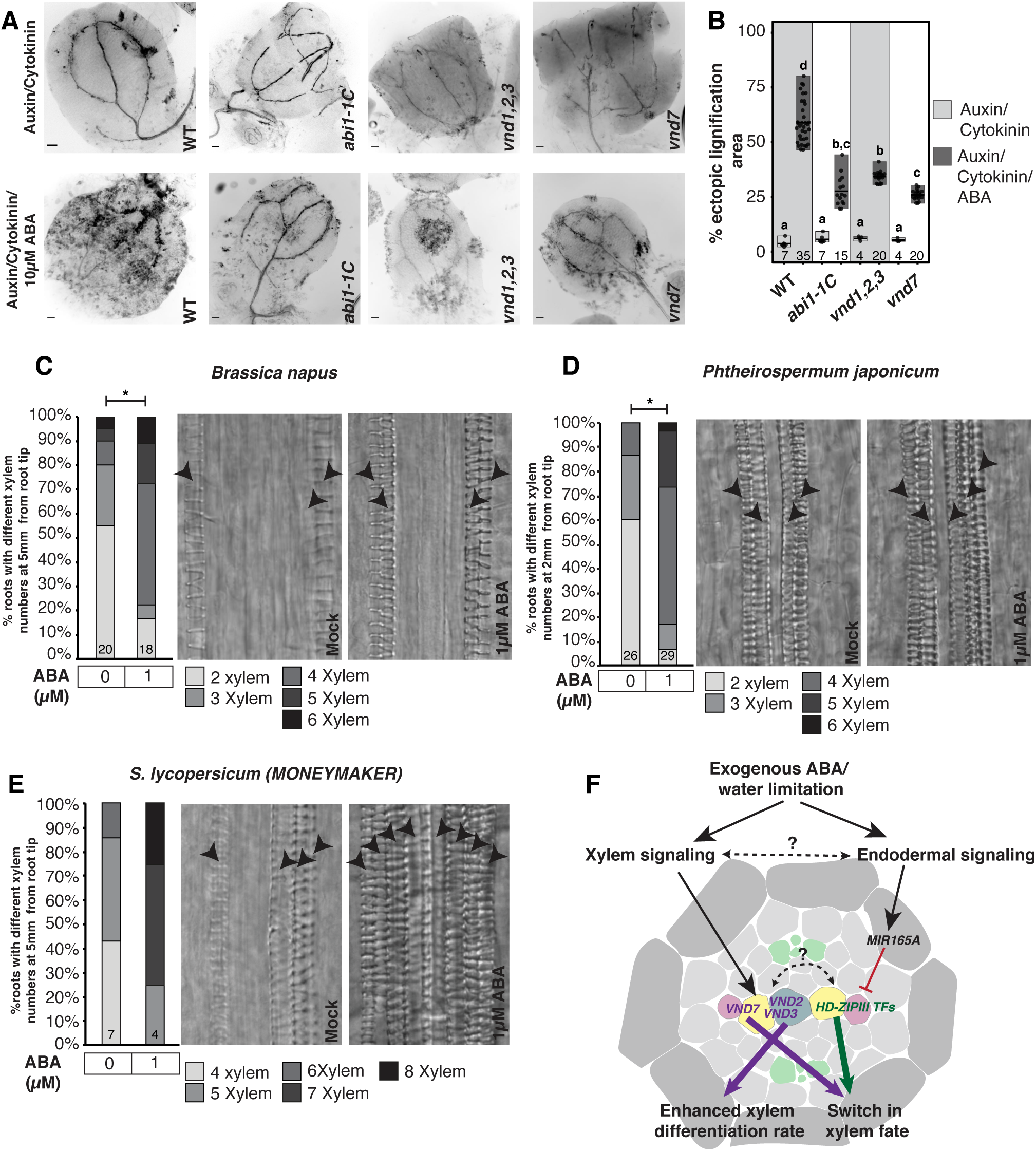
ABA induces ectopic lignification in Arabidopsis cotyledons and promotes xylem differentiation in several eudicot species. **(A)** Fluorescent micrographs showing the formation of ectopic lignification in wild type (WT), *abi1-1C, vnd1 vnd2 vnd3* (*vnd1,2,3)* and *vnd7* cotyledons after *in vitro* culture in auxin-cytokinin containing media with or without ABA. Ectopic lignification is visualized using lignin autofluorescence, and appear as dark spots on the cotyledons. **(B)** Quantification of ectopic lignification area in *in vitro* cultured WT, *abi1-1C, vnd1,2,3* and *vnd7* cotyledons. **(C-E)** Quantification of total number of lignified xylem vessels at specific distances from the root tip in *Brassica napus* (C), *Phtheirospermum japonicum* (D) and *Solanum lycopersicum (Money Maker)* (E) after mock and 1µM ABA treatment accompanied by representative images. **(F)** Model showing genetic components regulated cell autonomously by ABA to mediate two different phenotypic effects. The ABA signaling in the stele activates *VND2, VND3* and *VND7.* While *VND2* and *VND3* are mainly involved in ABA mediated enhancement of xylem differentiation rate, *VND7* mediates the switch in xylem morphology from pitted to a spiral or reticulate form. In the endodermis, ABA signaling activates a mobile miRNA, miR165 which results in a downregulation of stele expressed HD-ZIP III transcription factors to alter xylem fate [2,16]. Scale bars are 1mm in A. Statistics: * in C, D represent P<0.05, Fisher’s Exact test. In B, black dots represent individual measurements. Numbers at the bottom of the bars in B-E represent the number of individuals analyzed.

### ABA promotes xylem differentiation in several eudicot species

To understand the conservation in plant xylem responses upon stress, we analyzed the root xylem upon ABA treatment in five different eudicot species (Fig. 4C-E, S4D-F). We found that *Brassica napus* and *Brassica rapa* (Brassicales, Rosidae), *Nicotiana benthamiana* and *Solanum lycopersicum* (Solanales, Asteridae) and *Phtheirospermum japonicum* (Lamiales, Asteridae) all displayed early xylem differentiation and a higher number of xylem strands compared to mock condition, similar to what we had observed in Arabidopsis (Fig.4C-E,S4D-F). Furthermore, we observed a significant upregulation of putative tomato *VND1 (Solyc02g083450), VND4 (Solyco08g079120)* and *VND6* (Solyc06g065410) homologs after 6h of 1µM ABA treatment (Fig.S4G). These results suggest a conservation in molecular and phenotypic responses to ABA among eudicots.

Taken together, here we provide insights into the molecular regulation underlying the observed xylem developmental plasticity in Arabidopsis, by showing that ABA signaling triggers alterations in xylem cell developmental trajectories, both affecting their fate and their rate of differentiation, through the activation of distinct transcriptional regulators belonging to the same gene family, the VND genes (Fig. 4F). Several pieces of evidence indicate that ABA signaling acts within the xylem cells to trigger the activation of these factors to accomplish these feats. However, ABA also acts non-cell autonomously via the activation of miR165 in the endodermis, reducing levels of HD-ZIP III transcription factors in the stele (Fig. 4F) [2,16]. Intriguingly, both pathways appear important for the determination of xylem cell fate. While studies of gene regulatory networks have uncovered a complex interplay of VND and HD-ZIP III transcription factors [19], it remains to be seen how these factors temporally may interplay within the pluripotent xylem precursor cells to determine xylem cell fate, under normal growth conditions and during stress.

The two distinct phenotypic changes observed under ABA treatment and water limiting conditions may contribute two distinct advantages to the plant. A change towards more protoxylem-like cells may provide improved resistance to embolism that may form and affect water transport during conditions of reduced water availability [22]. Early formation of metaxylem, on the other hand, has been shown to enhance hydraulic conductance and increase drought resistance [6]. A subset of Arabidopsis accessions constantly displays early metaxylem development, however, this is associated with enhanced pathogen sensitivity, likely explaining why other accessions instead display this feature as a plastic trait in response to ABA-mediated stresses. Furthermore, a recent study identified a maize mutant defective in a VND-homologue that displayed symptoms of water stress under normal conditions due to defective protoxylem cells in adult plants [23]. This suggests that VND transcription factor-dependent xylem cell acclimation to stress is a trait whose evolution preceded the divergence of monocots and eudicots, and thus unite most angiosperms. The ABA-VND regulation may thus be a potentially universal molecular toolkit for xylem cell developmental adjustments with utility for breeding of drought resilient crop plants.

## Acknowledgements

We thank J. R. Dinneny, Stanford University, (ABA signaling reporter lines and *abi1-1* driver lines), T. Demura, NAIST, (VND mutant and reporter lines), M. Englund for technical assistance, and Nottingham Arabidopsis Stock Centre for seeds. This study was supported by funding from Nilsson Ehle Foundation, Lars Hiertas Minne, Lundell PO scholarship (to P.R.), a Wallenberg Academy Fellowship (KAW2016.0274 to C.W.M.), Vetenskapsrådet (to C.W.M and S.M.), and FORMAS (2017-00857 to A.C.).

## Author contributions

Conceptualization, P.R., A.C. and F.A.; Investigation, P.R., F.A., S.M. and V.N.; Writing – Original Draft, P.R.; Writing – Review & Editing, P.R., A.C., C.W.M. and F.A.; Funding acquisition, P.R., A.C. and C.W.M.; Supervision, A.C., E.A.M. and C.W.M.

## Supplementary material

**Figure S1: ABA affects both xylem differentiation fate and rate in Arabidopsis roots (A)** Quantification of root lengths in mock and ABA treated WT roots. **(B)** Representative DIC images of xylem morphological changes in 1µM ABA treated WT roots quantified in Figure 1E. **(C-D)** Quantification of xylem morphology (C) and *imx* differentiation rate (D) changes in ABA treated roots after transfer and growth for two days in mock, M, or ABA, A conditions and further transfer for growth for another two days under mock or ABA conditions. **(E)** Representative DIC images showing the xylem pattern after transfer of ABA treated roots to ABA or mock plates. **(F)** Quantification of xylem morphology changes in WT and *abi1-1 C* after 48h 1µM ABA treatment. **(G-H)** Quantification of xylem morphology (G) and *imx* differentiation rate (H) changes in *pSCR:abi1-1* lines after 48h 1µM ABA treatment. **(I-J)** Confocal micrograph showing ABA response domain after ABA treatment visualized using *6xABRE_R:erGFP* reporter (I) and after PEG treatment in *6XABRE_A:erGFP* (J). Radial optical sections were taken at 20 and 60µm from the QC in I and J. Magenta: Propidium iodide, Green: GFP and white arrow heads: xylem axis. RX, reticulate xylem; PX, protoxylem; *px*, protoxylem position; *omx*, outer metaxylem position; *imx*, inner metaxylem position. in C, F and G. Scale bars: 50µm in B, E, I and J. Statistics: * in C, D, F, G represent P<0.05, Fisher’s Exact test. Numbers at the bottom of the bars in C, D, F, G and H represent the number of individuals analyzed.

**Figure S2: ABA signalling within the xylem activates VND transcription factors (A)** Activity domains of different enhancer trap lines used for transactivation of *abi1-1* under mock or ABA treatment. **(B)** Quantification of xylem morphology changes observed in *abi1-1* transactivation lines. Numbers at the bottom of the bars in B represent the number of individuals analyzed. **(C)** DIC images showing xylem breaks observed in *J0571>>abi1-1* lines. **(D-F)** Expression of xylem development genes as determined by qRT-PCR after 4h (D) and 8h (E) 1µM ABA treatment in 1mm root tips (D,E) and 50µM ABA treatment for 4h in whole roots (F). **(G)** Confocal images showing transcription domains of *VND3* and *VND7* within and above the meristematic zone. **(H)** Confocal images showing the activation of *pVND3::YFP-NL*S reporter after 1µM ABA treatment for 6-8h. **(I)** Quantification of YFP nuclear intensity in mock and ABA treated *pVND3::YFP-NL*S. Scale bars: 50µm in A, C, H and I. Statistics: * in B, represent P<0.05, Fisher’s Exact test; * in D-F and I represent P<0.05, two tailed Student’s t-test. In D-F, all values are normalized to the average of mock treated samples. Black dots in D-F represent biological replicates and grey dots represents each quantified nucleus in I.

**Figure S3: VNDs regulate plasticity of xylem fate and differentiation rate (A-B).** Quantification of xylem morphology (A) and differentiation in inner metaxylem position (*imx*) at 7 mm from the root tip (B) after mock and 1µM ABA treatment (48h) in wild type (WT), single and double *vnd* mutants. **(C)** Quantification of xylem differentiation observed after polyethylene glycol (PEG) treatment in WT, *vnd1 vnd3* (*vnd1,3*) and *vnd2 vnd3* (*vnd2,3*) mutants. Plants were categorized depending on the distances at which the first lignified *imx* xylem vessel was observed. **(D)** Quantification of root lengths in *vnd1,3, vnd2,3, vnd7* and (*vnd1 vnd2 vnd3*) *vnd1,2,3* mutants after PEG treatment. **(E)** Venn diagram illustrating the VND dependence of several ABA regulated xylem enriched genes [11]. Genes were considered to be VND dependent if they were significantly differentially expressed in WT upon ABA treatment but not in *vnd1,2,3* or *vnd7* mutants. **(F)** Quantification of distance from root tip to a lignified xylem vessel in the outer metaxylem position (*omx*) in WT, *vnd2,3, vnd1,2,3* and *vnd7* mutants. **(G)** Quantification of distance from root tip to a lignified xylem vessel in *imx* in WT, *vnd1,3, vnd2,3* and *vnd7* mutants. **(H)** Quantification of xylem morphology changes in *vnd2 vnd3 vnd7* (*vnd2,3,7*) and *vnd1 vnd2 vnd3 vnd7* (*vnd1,2,3,7*) mutants. Statistics: * in A, B, C represent P<0.05, Fisher’s Exact test. In D and F-G, *a,b,c,d* represent groups with significant differences, one-way ANOVA with Tukey’s post-hoc testing (P<0.05). Black dots in D, F-G represent biological replicates. Numbers at the bottom of the bars in A-D and F-H represent the number of roots analyzed.

**Figure S4: ABA induces ectopic lignification in Arabidopsis cotyledons and promotes xylem differentiation in several eudicot species. (A)** Venn diagram of differentially expressed genes (DEG) upon ABA treatment and upon trans-differentiation of xylem cells in Arabidopsis cell suspension cultures [12]. Genes were considered to be VND dependent if they were significantly differentially expressed in WT upon ABA treatment but not in *vnd1 vnd2 vnd3* (*vnd1,2,3*) or *vnd7* mutants. **(B)** Zoomed in fluorescent micrographs showing autofluorescence of lignified cells in the cotyledons subjected to auxin/cytokinin/ABA treatment. **(C)** Ectopic lignification in WT cotyledons treated with auxin/cytokinin/ABA visualized using Basic Fuchsin staining. **(D-F)** Quantification of total number of lignified xylem vessels at specified distances from the root tip in *B. rapa* (D), *N. benthamiana* (E) and *S. lycopersicum (Tiny Tim)* (F) after mock and 1µM ABA treatment accompanied by representative images showing an increase in xylem number. **(G)** qRT-PCR of *VND* homologs in tomato roots after 1µM ABA treatment for 6h. Scale bars are 1mm in B and C. Statistics: * in E represent P<0.05, Fisher’s Exact test; * in H represent P<0.05, two tailed Student’s t-test. Numbers at the bottom of the bars in D-F represent the number of roots analyzed.

**Table S1. RNA sequencing analysis of Col-0, *vnd1 vnd2 vnd3* and *vnd7* mutants under mock or ABA treatment.**

**Table S2. List of primer sequences used in this study**

## STAR methods

### Plant material and growth conditions

Plant material used was *Arabidopsis thaliana, Nicotiana benthamiana, Phtheirospermum japonicum* and *Solanum lycopersicum* (cv. Money maker and Tiny Tim). Seeds were surface sterilized using 70% Ethanol for 20 mins and 95% Ethanol for 2-3 mins, and then rinsed in water four times. The seeds were imbibed and stratified for 48h at 4°C, and plated on 0.5xMS medium with 1% Bactoagar. For all experiments, plants were grown vertically on 25µm pore Sefar Nitex 03-25/19 mesh, and transferred to new plates by transferring the mesh with the plants on for minimal disturbance. For experiments involving transfer from ABA back to mock conditions, seedlings were instead transferred individually to prevent effects of residual ABA on the mesh. For ABA (Sigma) treatment, stock solutions of 50mM and 5mM ABA in 95% ethanol were used to make plates with ABA concentrations as indicated. Treatment with polyethylene glycol, was done with PEG 8000, as previously described [1,32].

All plant growth was carried out in long day conditions, 16h light (22°C) and 8h darkness (20°C) at light intensity of 110µE/m^2^/s. For Arabidopsis phenotyping experiments, two-day old seedlings were transferred to 1µM ABA containing plates for treatments of times indicated. For gene expression analysis, 4-5-day old seedlings were used. For phenotyping other species, seedlings were grown until roots reached approximately 1cm in length before transfer to ABA-containing plates.

All mutants or lines used in this study were in Col-0 background. Mutants and transgenic lines used in this study include *abi1-1C* [24], *pSCR:abi1-1* [10], *UAS::abi1-1* [10], *vnd* mutants (*vnd1, vnd2, vnd3, vnd1 vnd2, vnd2 vnd3, vnd1 vnd3, vnd1 vnd2 vnd3, vnd6, vnd7*) [25]. For generation of the *vnd1 vnd2 vnd3 vnd7* quadruple mutant, the *vnd1 vnd2 vnd3* triple mutant was crossed to *vnd7* mutant, and segregating F2 seedlings where genotyped using the primers listed in Table S1. The ABA responsive reporters used in this study are from [9] and the VND transcriptional reporters are from [12]. For tissue specific expression of *abi1-1, UAS::abi1-1* were crossed to Haseloff driver lines [26] and the resulting F1 seedlings were used for further analysis.

### Phenotypic analysis

#### Xylem morphology quantification

For analysis of xylem morphological changes, roots were mounted directly in chloralhydrate solution and visualized as described previously [2]. For quantification of phenotypes, the entire primary root or part of the root grown during the treatment times were analyzed for differences from wildtype pattern, separately for the distinct xylem axis positions (*px, omx* and *imx*). Phenotypes were categorized and the number of plants displaying a certain phenotype was used to calculate the frequency. Presence of more than one phenotype occurring in the same root was classified into a separate category.

### Determination of point of xylem differentiation initiation

For determination of the point of xylem differentiation initiation, i.e. where secondary cell wall and lignification can be detected first relative the root tip, roots were cleared and stained with ClearSee solution containing calcofluor white and basic fuchsin [27]. Tile scans of roots from the root tip was acquired using Zeiss LSM780 inverted Axio Observer with supersensitive GaAsP detectors. Distances from the root tip to xylem vessel with bright fuchsin staining (lignin) at different positions in the xylem axis was measured by drawing a line from the root tip to the point of lignification using Zeiss Zen software.

### Xylem differentiation at the *inner metaxylem* position

For early *imx* differentiation phenotypes, roots were mounted in chloralhydrate solution parallel to each other with root tips aligned on glass slides. A line was draw on the glass slide at a distance of 7mm from the root tip and this 7mm section of the root from the root tip was analyzed for the presence of lignified metaxylem. Roots were scored for presence or absence of a lignified *imx* using Zeiss Axioscope A1 microscope. For *B. napus, B. rapa* and *S. lycopersicum*, roots were mounted similarly to Arabidopsis and the number of xylem vessels at 5mm from the root tip was quantified. For *P. japonicum* and *N. benthamiana*, xylem vessel number was quantified at 2mm from the root tip.

The number of primary roots analyzed in each experiment is represented in the individual figures. Most experiments were repeated at least three times with similar results.

### Confocal analysis

Roots were mounted in 40µM propidium iodide (PI) solution between two cover slips and imaged immediately. Confocal micrographs were captured using Zeiss LSM780 inverted Axio Observer with supersensitive GaAsP detectors. For calcofluor white 405nm laser was used for excitation and emission wavelengths 410-524nm were captured in the detector. For basic fuchsin, 561nm excitation; 571-695nm emission. For reporter lines expressing GFP and stained with PI: 561nm excitation and 650-719, emission for PI and 488nm excitation and 500-553nm emission for GFP. For reporter lines expressing YFP, 514nm excitation for both YFP and PI, 518-562 emission for YFP and 651-688nm emission for PI was used. For experiments involving quantification of fluorescence intensity all imaging parameters were kept the same when imaging mock and ABA-treated roots. The Zeiss Zen software was used to quantify YFP intensity. Region of Interests (ROI) encompassing nuclei in the Arabidopsis root meristem were used to measure average fluorescence intensity. Nuclei from similar regions in the root was used for mock and ABA treated samples.

### Xylem trans-differentiation of cotyledon cells

For vascular induction in Arabidopsis cotyledons we followed the protocol used for xylem induction in cotyledons using bikinin with minor modifications [21]. The modifications include the following: 1. In the induction medium, all components were like in [21] except that bikinin was replaced with 10µM ABA. 2. The time for induction was increased from 4 days to 6 days. At the end of the 6-day induction period, cotyledons were fixed and cleared before visualization of autofluorescence upon UV exposure. The area of ectopic lignification (autofluorescence) was calculated using Image J and normalized to the total cotyledon area. Cotyledon veins were excluded from the quantification.

### Expression analysis by quantitative RT-PCR

qRT-PCR analysis was performed as previously described [2]. For Arabidopsis, either 1mm root tips or whole roots were used. For *S. lycopersicum* (cv Tiny Tim), whole roots were collected after 1µM ABA treated for 6h. Primers used in this study are listed in Table S2. Three biological replicates were used for all samples and individual data points are represented in graphs. APT1 and GAPDH for Arabidopsis and ACTIN and TIP41 for tomato was used as reference genes, respectively.

### RNAseq analysis

Five day old Arabidopsis seedlings of Col-0, *vnd1 vnd2 vnd3* and *vnd7* were treated for 8h with 1µM ABA or mock, respectively. The lower part of the root (1 cm) was collected, RNA was extracted with Qiagen RNeasy Plant Mini Kit, and sequenced by Novogene on their Illumina sequencing platform with paired-end read length of 150 and 250-300bp cDNA library. Mapping and differential expression analysis were done as follows. Briefly, mapping to the reference genome was done using Hisat2, count files generated using HTSeq and differential expression analysis was done using DESeq2. Comparison of mock and ABA-treated samples of the respective genotype and comparing WT mock samples with the *vnd1 vnd2 vnd3* mock and *vnd7* mock samples were used to identify differentially regulated genes. Genes with an adjusted p-value < 0.05 were considered differentially expressed. Genes were annotated as VND dependent if they were differentially expressed in Col-0 upon ABA treatment but not in the *vnd* mutants. Genes were annotated as VND independent if they were differentially expressed both in Col-0 and in the *vnd* mutants upon ABA treatment.

### Statistical analysis

For categorical data, Fisher’s exact test was used and p-values less than 0.05 were considered significant. For other data, One-way Anova or Student’s t-test was used. Statistical tests and significance threshold used are mentioned in the figure legends.

