## Supplementary Figures 1-4 for "Bipartite influence of abscisic acid on xylem differentiation trajectories is dependent on distinct VND transcription factors in Arabidopsis"

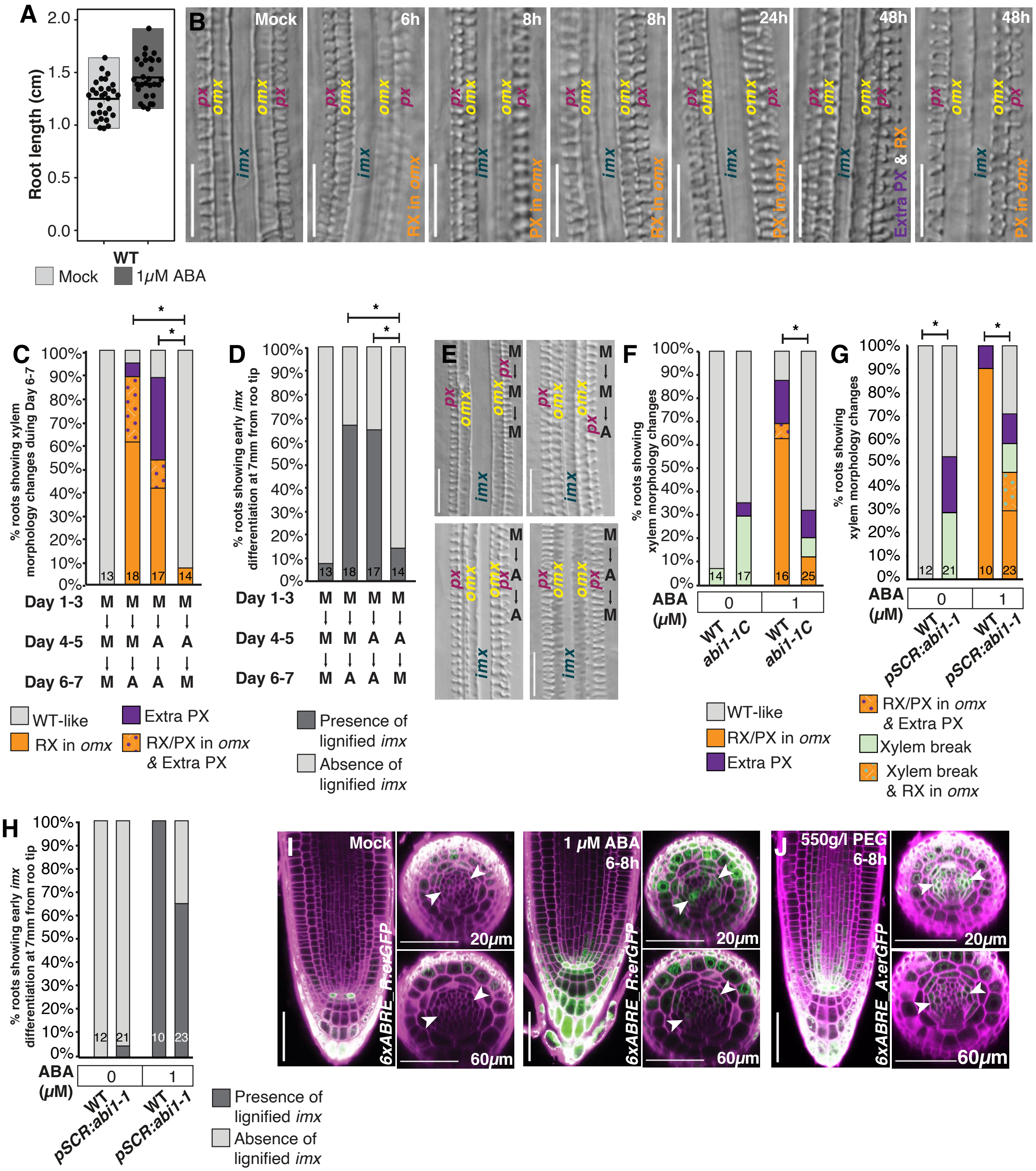


**Figure S1: ABA affects both xylem differentiation fate and rate in Arabidopsis roots**

**Figure S1: ABA affects both xylem differentiation fate and rate in Arabidopsis roots**

**(A)** Quantification of root lengths in mock and ABA treated WT roots. **(B)** Representative DIC images of xylem morphological changes in 1µM ABA treated WT roots quantified in Figure 1E. **(C-D)** Quantification of xylem morphology (C) and *imx* differentiation rate (D) changes in ABA treated roots after transfer and growth for two days in mock, M, or ABA, A conditions and further transfer for growth for another two days under mock or ABA conditions. **(E)** Representative DIC images showing the xylem pattern after transfer of ABA treated roots to ABA or mock plates. **(F)** Quantification of xylem morphology changes in WT and *abi1-1 C* after 48h 1µM ABA treatment. **(G-H)** Quantification of xylem morphology (G) and *imx* differentiation rate (H) changes in *pSCR:abi1-1* lines after 48h 1µM ABA treatment. **(I-J)** Confocal micrograph showing ABA response domain after ABA treatment visualized using *6xABRE_R:erGFP* reporter (I) and after PEG treatment in *6XABRE_A:erGFP* (J). Radial optical sections were taken at 20 and 60µm from the QC in I and J. Magenta: Propidium iodide, Green: GFP and white arrow heads: xylem axis. RX, reticulate xylem; PX, protoxylem; *px*, protoxylem position; *omx*, outer metaxylem position; *imx*, inner metaxylem position.  in C, F and G. Scale bars: 50µm in B, E, I and J.  Statistics: * in C, D, F, G represent $P<0.05$, Fisher's Exact test. Numbers at the bottom of the bars in C, D, F, G and H represent the number of individuals analyzed.

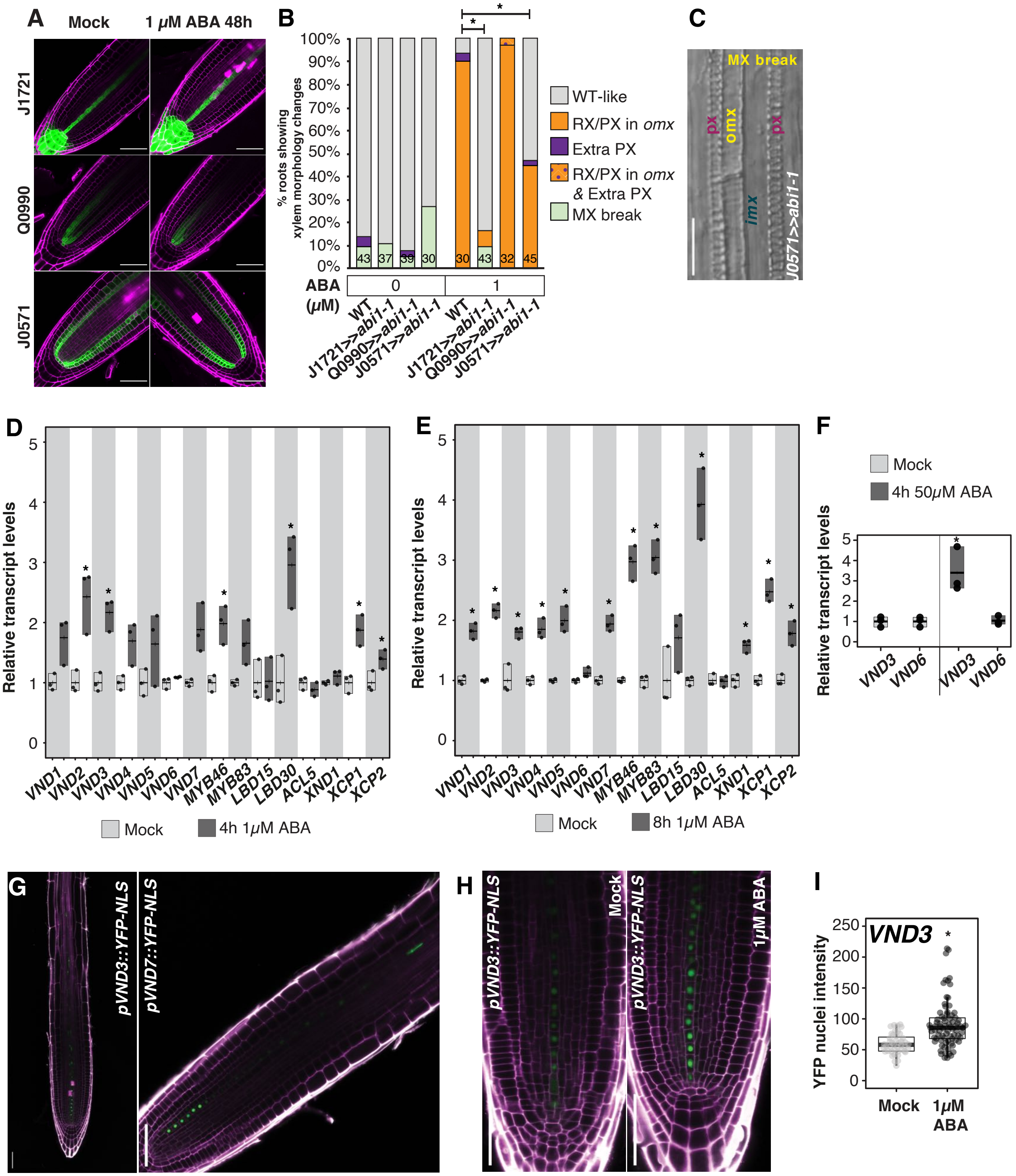


**Figure S2: ABA signalling within the xylem activates VND transcription factors**

**Figure S2: ABA signalling within the xylem activates VND transcription factors**

**(A)** Activity domains of different enhancer trap lines used for transactivation of *abi1-1* under mock or ABA treatment. **(B)** Quantification of xylem morphology changes observed in *abi1-1* transactivation lines. Numbers at the bottom of the bars in B represent the number of individuals analyzed. **(C)** DIC images showing xylem breaks observed in *J0571>>abi1-1* lines. **(D-F)** Expression of xylem development genes as determined by qRT-PCR after 4h (D) and 8h (E) 1µM ABA treatment in 1mm root tips (D,E) and 50µM ABA treatment for 4h in whole roots (F). **(G)** Confocal images showing transcription domains of *VND3* and *VND7* within and above the meristematic zone. **(H)** Confocal images showing the activation of *pVND3::YFP-NL*S reporter after 1µM ABA treatment for 6-8h. **(I)** Quantification of YFP nuclear intensity in mock and ABA treated *pVND3::YFP-NL*S. Scale bars: 50µm in A, C, H and I.

Statistics: * in B, represent $P<0.05$, Fisher's Exact test; * in D-F and I represent $P<0.05$, two tailed Student's t-test. In D-F, all values are normalized to the average of mock treated samples. Black dots in D-F represent biological replicates and grey dots represents each quantified nucleus in I.

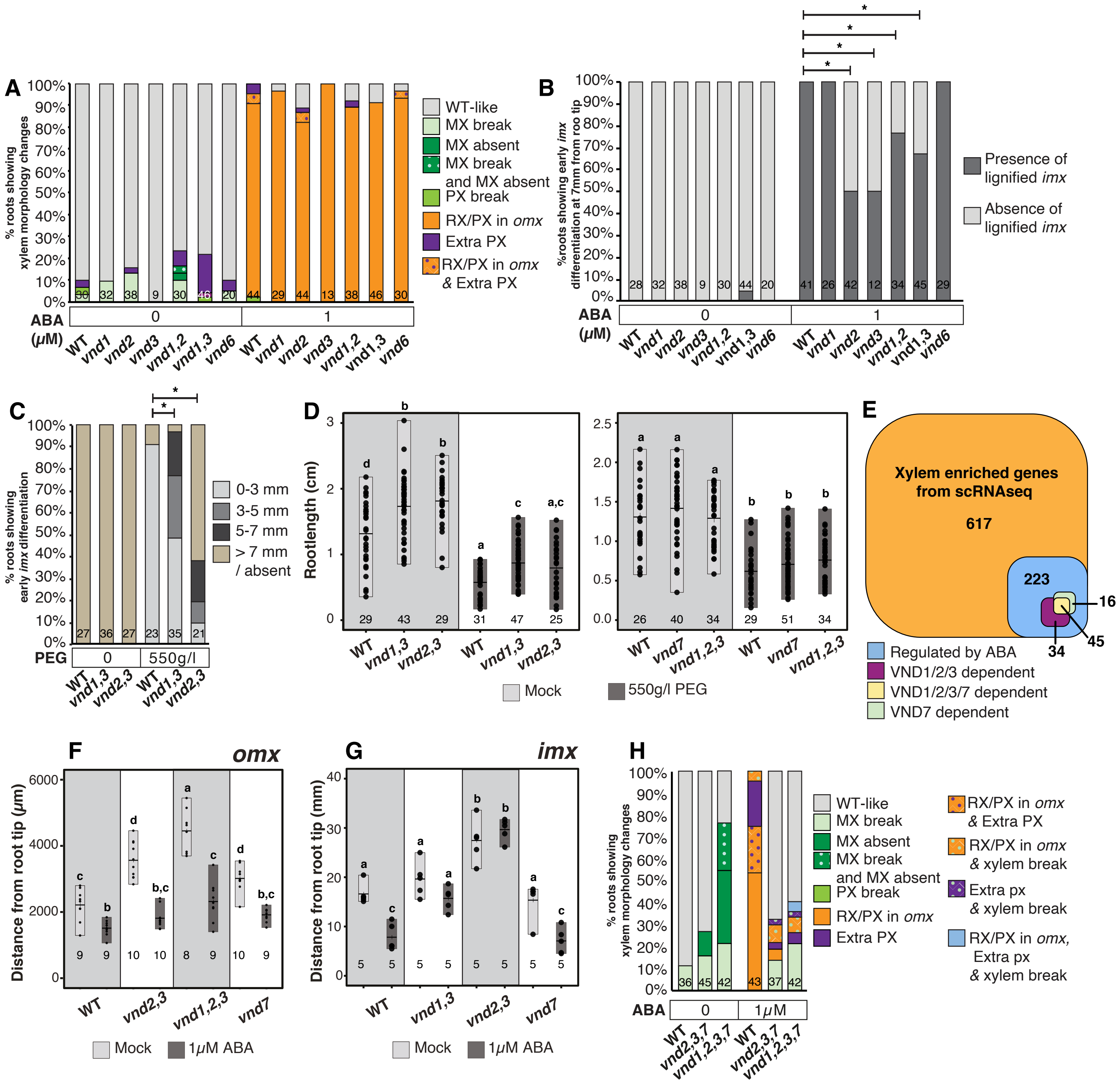


**Figure S3: VNDs regulate plasticity of xylem fate and differentiation rate**

**Figure S3: VNDs regulate plasticity of xylem fate and differentiation rate**

**(A-B).** Quantification of xylem morphology (A) and differentiation in inner metaxylem position (*imx*) at 7 mm from the root tip (B) after mock and 1µM ABA treatment (48h) in wild type (WT), single and double *vnd* mutants. **(C)** Quantification of xylem differentiation observed after polyethylene glycol (PEG) treatment in WT, *vnd1 vnd3* (*vnd1,3*) and *vnd2 vnd3* (*vnd2,3*) mutants. Plants were categorized depending on the distances at which the first lignified *imx* xylem vessel was observed. **(D)** Quantification of root lengths in *vnd1,3, vnd2,3, vnd7* and (*vnd1 vnd2 vnd3*) *vnd1,2,3* mutants after PEG treatment. **(E)** Venn diagram illustrating the VND dependence of several ABA regulated xylem enriched genes [11]. Genes were considered to be VND dependent if they were significantly differentially expressed in WT upon ABA treatment but not in *vnd1,2,3* or *vnd7* mutants. **(F)** Quantification of distance from root tip to a lignified xylem vessel in the outer metaxylem position (*omx*) in WT, *vnd2,3*, *vnd1,2,3* and *vnd7* mutants. **(G)** Quantification of distance from root tip to a lignified xylem vessel in *imx* in WT, *vnd1,3*, *vnd2,3* and *vnd7* mutants. **(H)** Quantification of xylem morphology changes in *vnd2 vnd3 vnd7* (*vnd2,3,7*) and *vnd1 vnd2 vnd3 vnd7* (*vnd1,2,3,7*) mutants. Statistics: * in A, B, C represent $P<0.05$, Fisher's Exact test. In D and F-G, *a,b,c,d* represent groups with significant differences, one-way ANOVA with Tukey's post-hoc testing ($P<0.05$). Black dots in D, F-G represent biological replicates. Numbers at the bottom of the bars in A-D and F-H represent the number of roots analyzed.

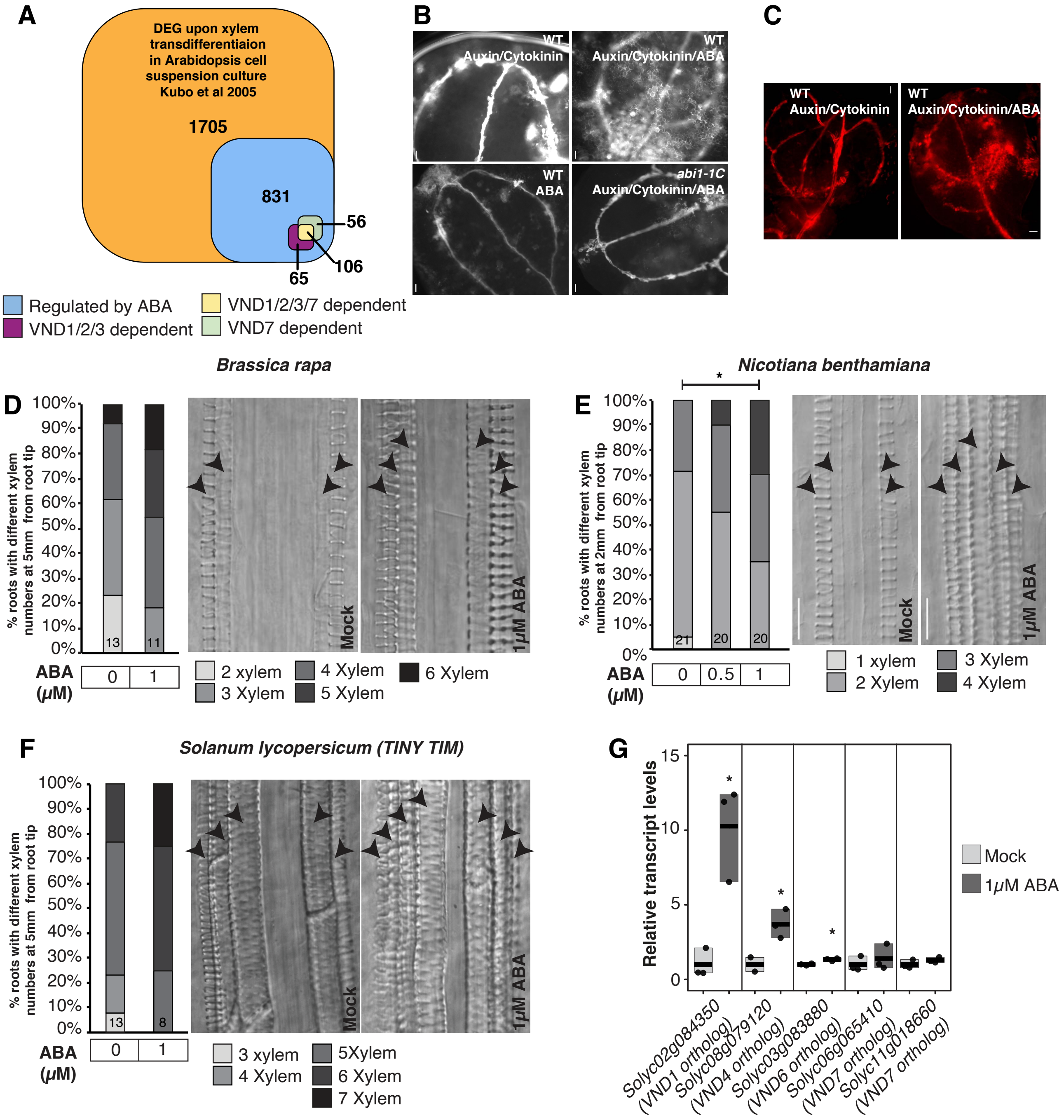


**Figure S4: ABA induces ectopic lignification in Arabidopsis cotyledons and promotes xylem differentiation in several eudicot species.**

**Figure S4: ABA induces ectopic lignification in Arabidopsis cotyledons and promotes xylem differentiation in several eudicot species.**

**(A)** Venn diagram of differentially expressed genes (DEG) upon ABA treatment and upon trans-differentiation of xylem cells in Arabidopsis cell suspension cultures [12]. Genes were considered to be VND dependent if they were significantly differentially expressed in WT upon ABA treatment but not in *vnd1 vnd2 vnd3* (*vnd1,2,3*) or *vnd7* mutants. **(B)** Zoomed in fluorescent micrographs showing autofluorescence of lignified cells in the cotyledons subjected to auxin/cytokinin/ABA treatment. **(C)** Ectopic lignification in WT cotyledons treated with auxin/cytokinin/ABA visualized using Basic Fuchsin staining. **(D-F)** Quantification of total number of lignified xylem vessels at specified distances from the root tip in *B. rapa* (D), *N. benthamiana* (E) and *S. lycopersicum (Tiny Tim)* (F) after mock and 1µM ABA treatment accompanied by representative images showing an increase in xylem number. **(G)** qRT-PCR of *VND* homologs in tomato roots after 1µM ABA treatment for 6h. Scale bars are 1mm in B and C. Statistics: * in E represent $P<0.05$, Fisher's Exact test; * in H represent $P<0.05$, two tailed Student's t-test. Numbers at the bottom of the bars in D-F represent the number of roots analyzed.
